# Differential Assembly of Native ENaC Complexes Across Mouse Epithelial Tissues

**DOI:** 10.64898/2026.01.23.701393

**Authors:** Arpita Bharadwaj, Joshua Curry, Xiao-Tong Su, Romina Barria Maturana, James A. McCormick, David H. Ellison, Isabelle Baconguis

## Abstract

The epithelial sodium channel (ENaC) governs sodium and fluid absorption in the lung, kidney, and colon, but the organization of native ENaC complexes has remained difficult to define because of their low abundance and biochemical instability. To enable direct analysis of native assemblies, we generated a knock-in mouse in which the endogenous γ subunit is fused at its C terminus to mVenus, a 3C protease cleavage site, and a 3xFLAG epitope (ENaCγ-VF). The tag preserves physiological ENaC function, as ENaCγ-VF mice display normal electrolyte handling, benzamil affinity, and amiloride-sensitive Na⁺:K⁺ responses indistinguishable from wild-type animals. Using fluorescence-detection size-exclusion chromatography and single-molecule pull-down, we directly monitor intact native ENaC complexes from lung, kidney, and colon and uncover marked tissue-to-tissue differences in channel abundance and apparent complex size. Dual-color analysis with a fluorescent Fab against ENaCα marks fully assembled αβγ channels, while γ-based fluorescence reports the broader population of γ-containing assemblies. In combination, the ENaCγ-VF line provides a biochemical anchor for identifying regulatory and trafficking proteins that co-purify with native ENaC complexes. These data show that ENaC architecture *in vivo* is heterogeneous, and establish ENaCγ-VF mice as a platform for dissecting how epithelial environments shape ENaC assembly, composition, and regulation.

## Introduction

Communication across epithelial barriers depends critically on the controlled movement of sodium ions, a process in which the epithelial sodium channel (ENaC) plays a central and indispensable role. Situated at the apical membrane of epithelial cells, ENaC drives sodium entry that underlies extracellular fluid balance, blood pressure regulation, and airway surface hydration^1–4^. Its activity is required for efficient alveolar fluid clearance in the lung^5, 6^, sodium reabsorption in the distal nephron^1^, and electrolyte absorption in the colon^7^. Dysregulation of ENaC function contributes to diseases such as cystic fibrosis^8^ and pseudohypoaldosteronism^9–12^, as well as multifactorial disorders including pulmonary edema^13^ and hypertension^14–25^. Hypertension alone affects more than one billion individuals worldwide and remains the leading modifiable risk factor for cardiovascular disease^26–28^, chronic kidney disease^29^, and cognitive decline^30^. These clinical associations have long underscored the need to understand how ENaC is assembled, regulated, and stabilized within its native environment.

Before recombinant expression was feasible, early conceptual models of ENaC biology emerged largely from electrophysiological studies of transepithelial transport, which inferred the presence of a selective sodium channel from its amiloride sensitivity and ionic selectivity^31^. These foundational observations established the physiological role of ENaC long before its molecular identity was known. The subsequent cloning of the α, β, and γ subunits provided the first molecular description for the channel and revealed a family of homologous proteins with conserved topology across vertebrate epithelia^32–34^. Recombinant expression systems soon enabled functional dissection through mutagenesis, proteolytic activation studies, and single-channel recordings, offering high-resolution insights into ENaC architecture^35–37^, gating^38^, and inhibition^39^.

Yet these advances captured only a portion of ENaC biology. Within native tissues, ENaC expression is highly compartmentalized^40^, its maturation requires sequential cleavage by distinct proteases^39, 41^, and its surface abundance is dynamically regulated by a network of accessory proteins and ubiquitin ligases^1, 42–44^. Electrophysiological and transcriptomic analyses have yielded valuable perspectives, but neither directly resolves how ENaC subunits assemble into functional complexes, nor how regulatory factors shape the architecture and stability of those assemblies. As a result, the molecular organization of ENaC *in situ*, its subunit stoichiometry, tissue-specific variability, and regulatory context, remain only partially defined. A fundamental obstacle has been the absence of tools enabling direct detection, purification, and visualization of ENaC under non-denaturing conditions. ENaC is expressed at exceptionally low abundance, biochemically labile, and embedded within lipid environments that complicate solubilization^45–47^. Attempts to recover native ENaC have traditionally yielded insufficient quantities for systematic analysis, limiting the field to indirect methods or recombinant models that cannot fully replicate the physiological assembly landscape.

To address this longstanding bottleneck, we developed a genetically engineered knock-in mouse line (ENaCγ-VF) in which the endogenous γ subunit is fused at its C terminus to mVenus, followed by a 3C protease site and a 3xFLAG epitope. This multifunctional tag enables fluorescence-based detection, affinity purification, and gentle protease-mediated elution of intact ENaC complexes while preserving native expression levels, regulatory control, and tissue specificity. Here, we validate that the tagged γ subunit supports normal ENaC function *in vivo* and demonstrate that the ENaCγ-VF line is compatible with fluorescence-detection size-exclusion chromatography (FSEC)^48^ and single-molecule pull-down (SiMPull)^49^, permitting direct quantification of native ENaC assemblies isolated from lung, kidney, and colon. In parallel, we developed a fluorescent Fab fragment targeting ENaCα, enabling dual-color, subunit-resolved analysis of ENaC architecture in native lysates. These tools establish a robust platform for dissecting ENaC assembly, composition, and regulation directly in physiological contexts. Because ENaC activity must be tuned to the distinct transport demands of different epithelia, tissue-specific differences in subunit assembly and associations with regulatory partners are likely to shape Na^+^ transport capacity and hormonal responsiveness^1–4, 39, 41–44^.

## Results

### Generation of the ENaCγ-VF mouse line

To enable detection and purification of native ENaC complexes, we designed a knock-in mouse line in which the endogenous *Scnn1g* locus (encoding ENaCγ) was modified to incorporate a C-terminal mVenus^Q69M^ fluorophore, a 3C protease recognition site, and a 3xFLAG epitope^50–52^ (Fig. 1a). This multifunctional design enables direct fluorescence detection, native-state affinity isolation, and protease-mediated elution of intact ENaC assemblies. The ENaCγ-VF allele was generated at the Jackson Laboratory using CRISPR/Cas9 to insert the tag in-frame at the endogenous *Scnn1g* stop codon^53, 54^. Initial characterization was performed in heterozygous ENaCγ-VF^+^/^-^ mice to assess whether C-terminal tagging of γENaC perturbed channel expression, trafficking, or subcellular localization in the presence of an unmodified endogenous allele. Heterozygous ENaCγ-VF^+^/^-^ mice exhibited normal appearance and growth, and immunostaining of kidney sections confirmed correct apical localization of the tagged γ subunit within the distal nephron (Fig. S1), consistent with preserved trafficking and expression^40^.

**Figure 1.**
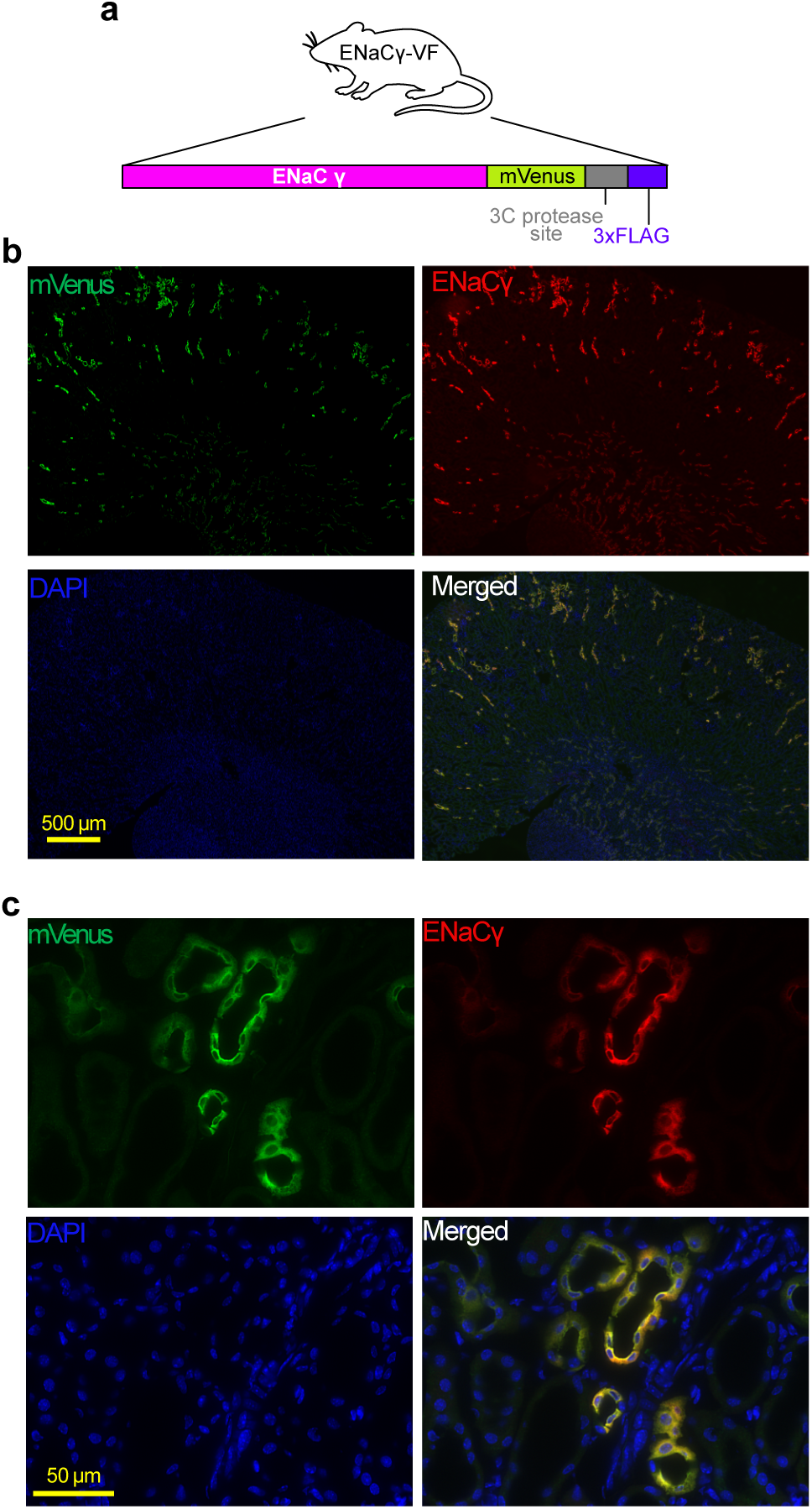
Immunofluorescence staining of kidney tissue from homozygous ENaCγ-VF mice. **a**. Schematic of the tagged ENaCγ subunit produced from the ENaCγ-VF allele, which includes a C-terminal mVenus, 3C protease site, and 3xFLAG tag. **b**. Representative immunofluorescence image showing co-localization of mVenus and ENaCγ in kidney tissue. mVenus signal was detected using a FITC-conjugated anti-GFP primary antibody (goat, Rockland, 1:100) and Alexa Fluor 488 donkey anti-goat secondary antibody. Endogenous ENaCγ was detected using a rabbit anti-ENaCγ primary antibody (StressMarq, 1:500) targeting the C-terminus, and Alexa Fluor 647 donkey anti-rabbit secondary antibody. DAPI is 4’,6-diamidino-2-phenylindole**. c**. Close-up view highlighting localization of tagged ENaCγ.

A central requirement for using this model to study native ENaC is that the modified γ subunit must independently support physiological channel function on its own. Because *Scnn1g* knockout is perinatally lethal^55^, viability in the homozygous state provides a stringent functional test of the tagged allele. Homozygous ENaCγ-VF/ENaCγ-VF mice were viable into adulthood, indicating that the mVenus-3C-3xFLAG fusion does not disrupt essential ENaC activity. These animals therefore served as the foundation for all subsequent physiological, biochemical, and biophysical analyses.

### Validation of the ENaCγ-VF mouse line

We validated the expression and localization of the tagged γ subunit in kidney tissue, where ENaC plays a well-defined role in sodium reabsorption. Immunostaining showed that mVenus-tagged γ was appropriately restricted to the distal nephron, consistent with the known distribution of native ENaC^40^ (Fig. 1b, c). Blood electrolyte levels in ENaCγ-VF mice were within the normal physiological range (Fig. 2a). To assess whether the tagged γ subunit supports normal channel function *in vivo*, we examined the response of ENaCγ-VF mice to amiloride, a well-characterized ENaC inhibitor^56^. As expected, pharmacological blockade increased the urinary Na⁺:K⁺ ratio, reflecting inhibition of ENaC-mediated sodium reabsorption and potassium secretion in the distal nephron. ENaCγ-VF mice exhibited an amiloride-induced increase in Na⁺:K⁺ ratio indistinguishable from wild-type controls (Fig. 2b), indicating preserved ENaC function. These findings demonstrate that homozygous ENaCγ-VF mice maintain normal electrolyte handling and are phenotypically comparable to wild-type littermates under basal conditions.

**Figure 2.**
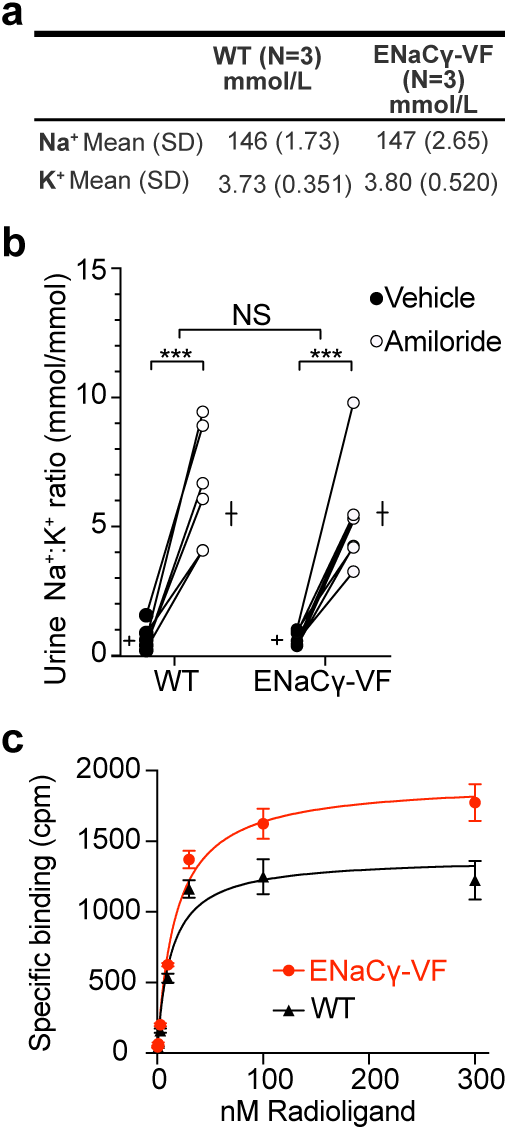
The ENaCγ-VF mouse line is physiologically indistinguishable from wild-type at baseline. **a**. Table showing plasma sodium (Na^+^) and potassium (K⁺) concentrations in wild-type C57BL/6 and homozygous ENaCγ-VF mice under baseline conditions. **b**. Amiloride response assay in wild-type C57BL/6 and ENaCγ-VF mice. The graph shows individual paired responses to vehicle and amiloride for each mouse, with overlaid bars indicating mean ± SEM (n = 6 per group). A two-way ANOVA revealed a significant effect of amiloride treatment compared with vehicle (***p < 0.0001), but no significant difference between genotypes (p = 0.4533), indicating that the tagged allele supports normal ENaC function. **c**. Saturation binding of [³H]benzamil to kidney membranes. Increasing concentrations of [³H]benzamil (0.5-300 nM) were incubated with kidney membranes for 2 hours min at 22 °C. Specific binding (circles) was determined by subtracting nonspecific binding measured in the presence of 100 µM unlabeled phenamil. Data represent one experiment performed in triplicate; points show mean ± SD of triplicate measurements. The curve was fit with a one-site specific binding model using nonlinear regression (GraphPad Prism) to obtain *K_d_*.

To complement these physiological measurements with a direct biochemical assessment of channel pharmacology, we performed radioligand filter-binding assays using tritiated [³H]-benzamil, a high-affinity ENaC antagonist^57^. Kidney membranes were selected for these experiments because renal ENaC activity underlies the *in vivo* amiloride challenge and provides a robust, well-defined tissue context in which ENaC-dependent sodium transport can be directly interrogated. Kidney membrane fractions were isolated, incubated with increasing concentrations of [³H]-benzamil, and filtered to separate receptor-bound from free radioligand. Specific benzamil binding was readily detectable in both wild-type and ENaCγ-VF membranes (Fig. 2c). The two genotypes exhibited comparable apparent dissociation constants (*K_d_* = 13 nM for wild-type and 17 nM for ENaCγ-VF). These values indicate that the C-terminal mVenus-3C-3xFLAG tag does not measurably alter benzamil affinity. The physiological and biochemical data establish that the tagged γ subunit preserves native ENaC function and pharmacological properties.

### Detection and characterization of native ENaC complexes using FSEC and SiMPull

The mVenus tag provides a sensitive fluorescence handle for tracking γ-containing ENaC assemblies in tissue lysates using both ensemble and single-molecule approaches (Fig. 3a). FSEC^48^ separates protein complexes by size under non-denaturing conditions and uses inline fluorescence to monitor tagged species as they elute from a calibrated gel-filtration column. Originally developed for recombinant membrane proteins, FSEC has proven effective for assessing complex formation, sample homogeneity, and expression levels^48, 58^. Here, we extend this approach to native tissues by using mVenus as an intrinsic reporter for γ-containing ENaC complexes.

**Figure 3.**
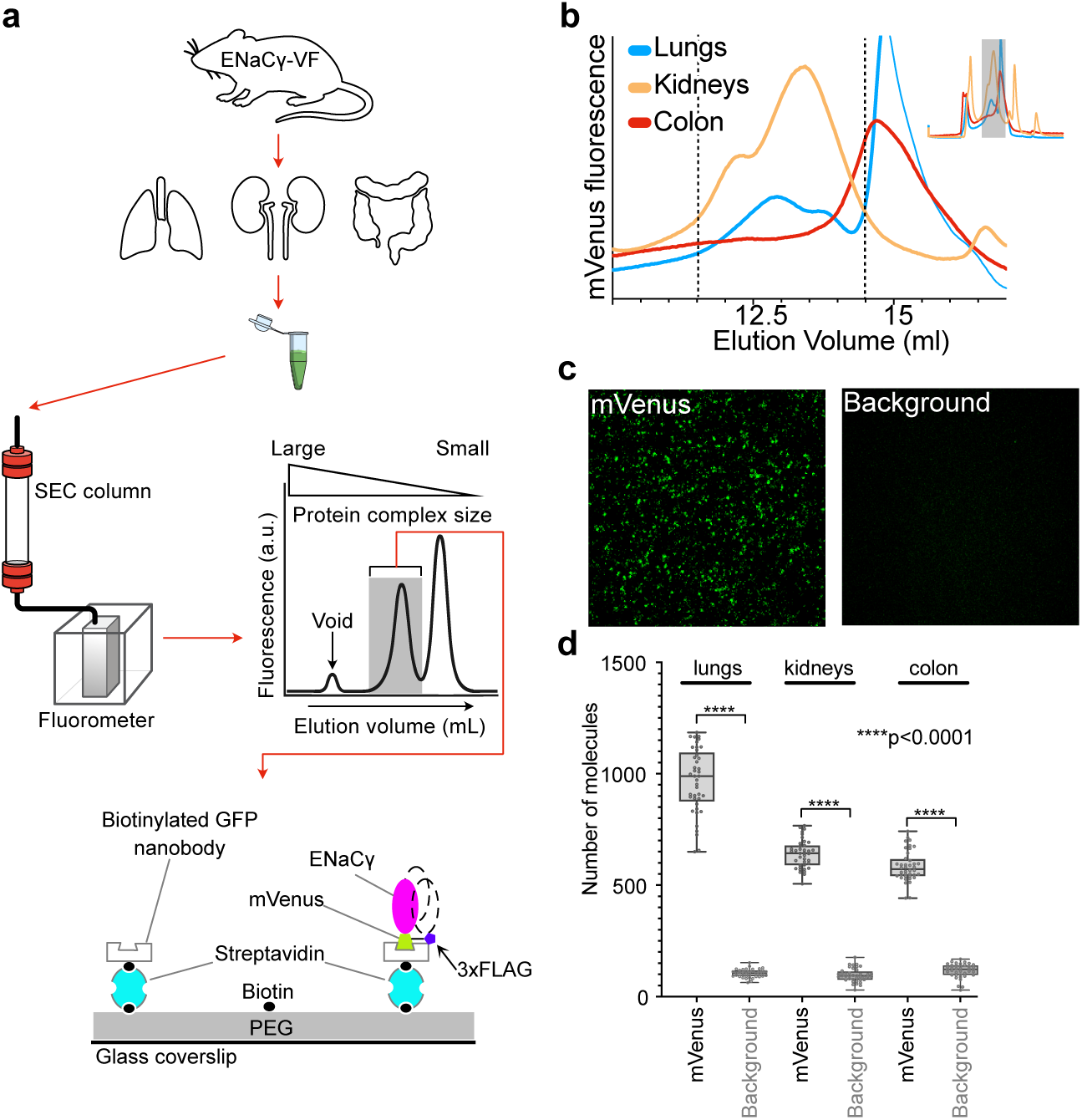
The ENaCγ-VF mouse enables subunit-specific detection of native ENaC complexes using FSEC and SiMPull. **a**. Schematic workflow illustrating the experimental pipeline. Tissues (lung, kidney, colon) were harvested from ENaCγ-VF mice, homogenized in TRIS-based buffer, and followed by solubilization. Cleared supernatants were injected onto a Superose 6 Increase column for FSEC analysis, and fractions corresponding to high-molecular-weight complexes were subsequently analyzed by SiMPull. **b**. Representative FSEC traces showing mVenus fluorescence from lung, kidney, and colon lysates. Fluorescent peaks indicate the presence of γ-containing ENaC complexes in each tissue. **c**. Representative TIRF images of SiMPull experiments from lung lysates. mVenus-tagged ENaC complexes were captured using a biotinylated anti-GFP/mVenus nanobody. A background control image was collected from a flow chamber lacking the nanobody to demonstrate specificity. **d**. Quantification of SiMPull signal from lung, kidney, and colon tissues. Each condition represents data pooled from 40 TIRF images per sample, analyzed using the ComDet plugin in ImageJ. Statistical analysis was performed in GraphPad Prism.

We applied FSEC to characterize native ENaC assemblies in lung, kidney, and colon tissues from ENaCγ-VF mice. Lysates were fractionated on a Superose 6 Increase column, and mVenus fluorescence was monitored across the elution profile (Fig. 3a). Lung and kidney lysates displayed well-defined fluorescence peaks eluting between 11 and 16 mL, consistent with the expected size range of multimeric ENaC complexes (Fig. 3b). In contrast, colon lysates produced substantially lower signal in initial FSEC runs. Free mVenus eluted at a later volume, allowing us to distinguish intact ENaCγ-containing complexes from unincorporated fluorophore and to verify sample integrity prior to downstream analyses.

To complement ensemble analyses and resolve the ENaC composition at the single-complex level, we integrated SiMPull^49^. SiMPull combines immunoaffinity capture with total internal reflection fluorescence (TIRF)^59^ microscopy to visualize individual protein complexes under non-denaturing conditions. This approach is well suited for low-abundance membrane proteins such as ENaC, offering attomole-scale sensitivity^60^ while preserving native subunit interactions. For selective capture of ENaCγ-VF, we employed a biotinylated anti-GFP/mVenus nanobody^61^ that binds to the mVenus fluorophore and enables immobilization of γ-containing assemblies on PEG-passivated, streptavidin-coated glass surfaces.

FSEC was used not only as an ensemble characterization but also as a preparative step to ensure that single-molecule measurements interrogated intact complexes. Fractions eluting before free mVenus, corresponding to high-molecular-weight γ-containing assemblies, were selected for SiMPull. Using this FSEC-guided workflow, robust mVenus signals were detected in lung and kidney samples, confirming efficient and specific capture of γ-containing complexes (Fig. 3c, d). For colon samples, where FSEC indicated lower ENaC abundance, lysates were concentrated prior to SiMPull, which enabled reliable single-molecule detection.

The combined use of FSEC and SiMPull provides a powerful, complementary platform for characterizing native ENaC assemblies. FSEC offers a rapid assessment of complex integrity and abundance, while SiMPull resolves individual complexes with single-molecule precision. These approaches establish a robust analytical platform for studying ENaC composition and assembly across multiple epithelial tissues. The observed tissue-dependent differences in ENaC abundance and apparent complex size are consistent with known variations in baseline Na^+^ transport and ENaC dependence across lung, kidney, and colon, and may contribute to how each epithelium matches channel activity to its specific absorptive demands^1–7, 40^.

### Dual-color FSEC using a fluorescent Fab against ENaCα

Having established that ENaCγ-VF provides a reliable fluorescence handle for tracking γ-containing assemblies across tissues, we next asked whether this platform could be extended to resolve the composition of native ENaC complexes at the subunit level. Because the mVenus tag reports specifically on the γ subunit, the analyses above define the behavior of γ-containing assemblies but do not directly address the incorporation of other ENaC subunits into the same complexes. A key advantage of the ENaCγ-VF allele is that it provides a stable biochemical anchor from which additional subunits can be assessed. To expand the detection capabilities of the system and enable subunit-resolved interrogation of native ENaC assemblies, we developed a fluorescent Fab directed against the extracellular domain of ENaCα.

The parent monoclonal antibody, 7B1, was originally raised against human ENaCα^35^, and sequence alignment revealed conservation between human and mouse α subunits within the predicted epitope (Fig. 4a, b), supporting the feasibility of cross-species recognition. To generate a monovalent, fluorescent reagent suitable for biochemical and biophysical studies, we cloned the variable regions of the 7B1 heavy and light chains into secretion-optimized expression constructs and fused the heavy chain C terminus to the red fluorescent protein mScarlet-I^62^, yielding 7B1-mScarlet (Fig. 4c). The Fab was secreted efficiently from HEK293 cells and isolated from conditioned media.

**Figure 4.**
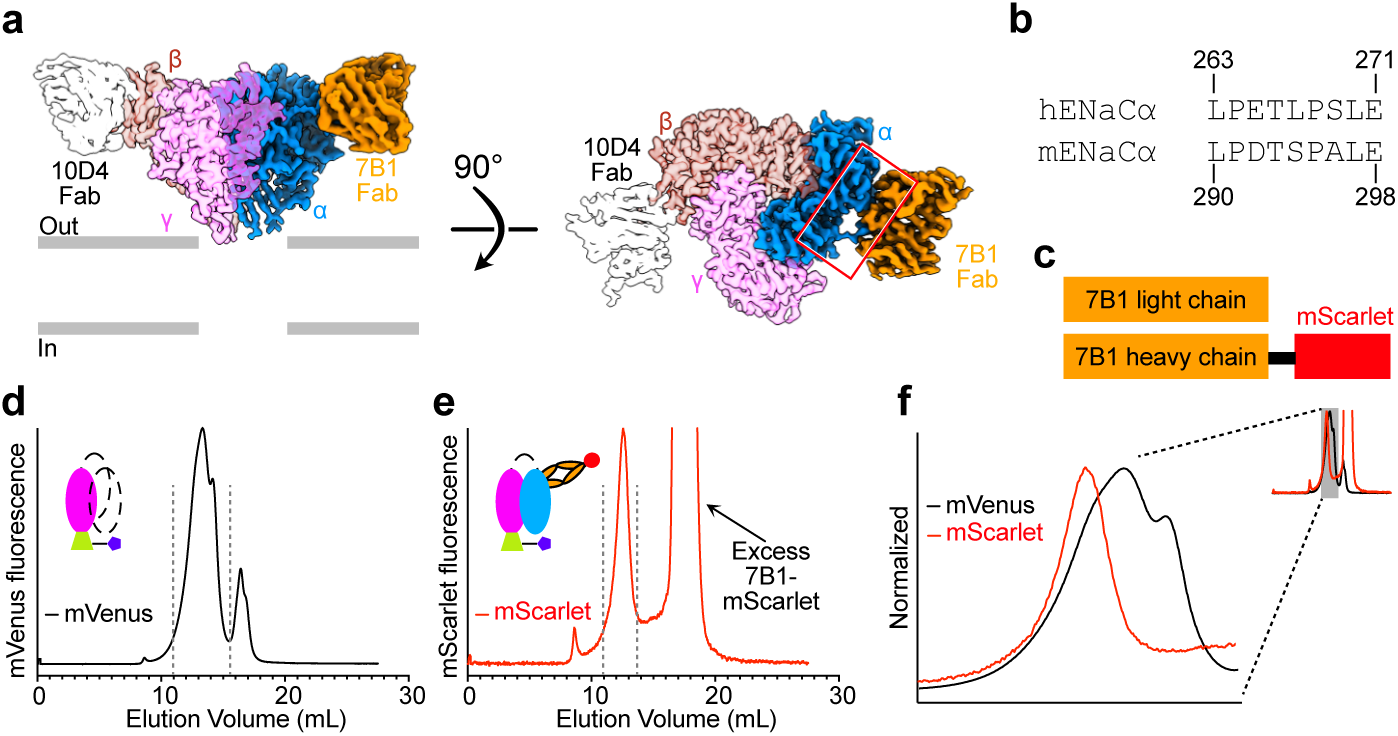
Dual-subunit detection of native ENaC complexes using 7B1-mScarlet and the ENaCγ-VF mouse line. **a.** Cryo-EM map of human ENaC in complex with two Fabs, 10D4 and 7B1, shown in side and top-down views. In the top-down view, a red box highlights the interaction site between 7B1 and the α subunit. **b.** Sequence alignment of the experimentally determined 7B1 epitope region in human and mouse ENaCα subunits, revealing high conservation and supporting the potential for cross-reactivity with mouse ENaC. **c**. Schematic representation of the 7B1-mScarlet Fab construct used to detect the ENaC α subunit. The variable regions of the 7B1 heavy and light chains were expressed as a Fab fragment in HEK293 cells, with mScarlet fused to the C-terminus of the heavy chain. **d**. FSEC trace monitoring mVenus fluorescence from lung lysates of ENaCγ-VF mice, showing the elution profile of γ-containing ENaC complexes. Dotted lines indicate the elution range for mVenus-tagged complexes (11-16 mL). **e**. FSEC trace from the same sample following 7B1-mScarlet fluorescence, revealing the elution profile of α-containing ENaC complexes. Dashed lines show that the α-associated signal is restricted to a narrower elution range (11-13 mL), suggesting that α-γ co-assemblies represent a more defined subset of the total γ-containing complexes. **f**. Overlay of the FSEC traces shown in panels **d** and **e**, plotted on the same axes to facilitate direct comparison of elution profiles and peak positions. The inset shows the full normalized traces for each condition. The main panel displays a magnified view of the gray-shaded region indicated in the inset, highlighting the relative alignment of the major peaks.

We first validated 7B1-mScarlet in a recombinant system expressing mouse ENaC composed of wild-type α and β subunits and a mVenus-tagged γ subunit, recapitulating the configuration of the ENaCγ-VF allele (Fig. S2a). Detergent-solubilized lysates were incubated with 7B1-mScarlet and analyzed by FSEC. mVenus and mScarlet fluorescence co-eluted in a single peak at the expected position for trimeric ENaC, whereas control samples lacking 7B1-mScarlet showed no mScarlet signal within this elution window (Fig. S2b). These results indicate that 7B1-mScarlet binds specifically to folded ENaC complexes containing the α subunit. To assess whether 7B1 recognizes α across different maturation states, we performed surface biotinylation of cells expressing recombinant mouse ENaC, lysed them, purified ENaC via the γ tag, and probed the resulting surface and intracellular fractions with 7B1-mScarlet (Fig. S2c); robust signal was detected in both biotinylated and non-biotinylated pools, indicating that the Fab can bind α in multiple maturation states rather than a single conformational end point (Fig. S2d).

We next asked whether this Fab could detect ENaCα within native γ-containing complexes isolated from tissue. Lung lysates from ENaCγ-VF mice were enriched for γ-containing assemblies by affinity capture through the C-terminal FLAG tag and eluted under non-denaturing conditions. Addition of 7B1-mScarlet to the eluted material followed by FSEC analysis revealed co-elution of mVenus and mScarlet fluorescence (Fig. 4d-f), demonstrating that α and γ subunits are incorporated into the same native complexes. Strikingly, while mVenus fluorescence extended across a broader range (11-16 mL), the 7B1-mScarlet signal was confined to the 11-13 mL window. These data establish dual-color FSEC as a non-denaturing approach for resolving the subunit composition of native γ-containing ENaC assemblies. The distinct behavior indicates that α and γ subunits partition into different subsets of γ-containing assemblies and highlights heterogeneity within the γ-containing population. The observations motivated further investigation into whether γ-containing assemblies of differing apparent sizes engage distinct regulatory protein complexes *in vivo*.

### ENaCγ-containing complexes associate with regulatory proteins *in vivo*

The heterogeneity revealed by dual-color FSEC raised the possibility that g-containing assemblies of different apparent sizes represent distinct molecular states. The presence of both a-containing complexes and broader g-containing assemblies is consistent with a model in which these complexes correspond to distinct maturation or stability checkpoints that tune ENaC expression^14–25, 39–44^. In this context, the 7B1-positive population represents α-containing complexes enriched within a narrower, higher molecular weight fraction, whereas γ-based fluorescence reports a continuum of γ-containing assemblies likely spanning multiple stages of folding, assembly, and regulatory interactions. To examine this possibility and extend our analysis beyond ENaC subunits alone, we turned to mass spectrometry to identify proteins that co-purify with γ-containing assemblies.

Large-scale affinity purification was performed from lung, kidney, and colon tissues of ENaCγ-VF mice using anti-FLAG enrichment followed by 3C protease cleavage to gently release native complexes under non-denaturing conditions (Fig. 5a). Fractions corresponding to the mVenus fluorescence peak were collected and subjected to mass spectrometry analysis (Fig. 5b). As expected, ENaCγ co-purified with ENaCα and β subunits (Fig. 5c), consistent with our dual-color FSEC data demonstrating α-γ co-assembly (Fig. 4f). Recovery of all three subunits confirms the integrity of the purified complexes and demonstrates that the ENaCγ-VF tag preserves native channel assembly.

**Figure 5.**
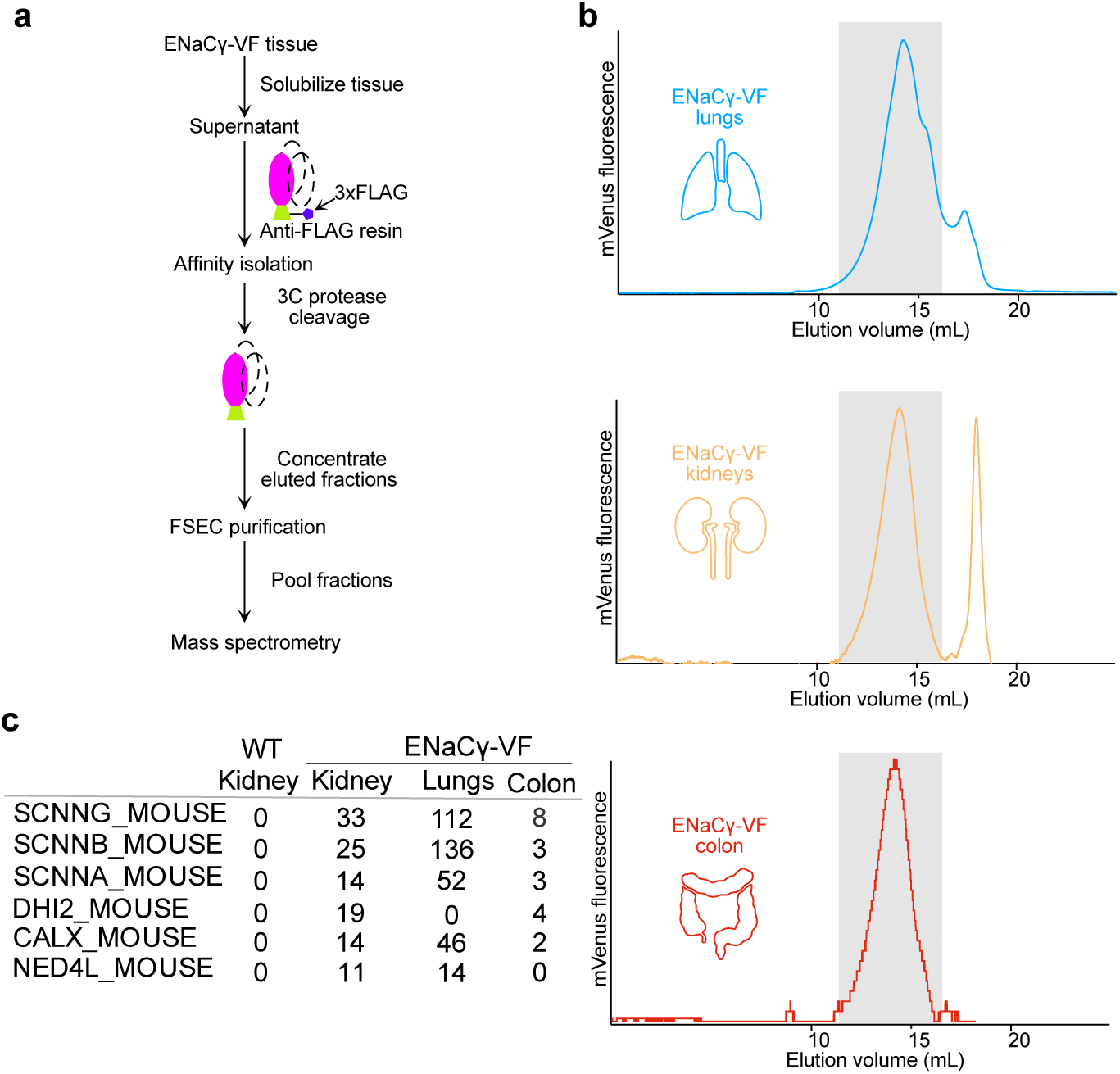
ENaC complex purification enables tissue-specific interactome analysis by mass spectrometry. **a.** Schematic workflow illustrating the purification pipeline. Tissues (lung, kidney, colon) from ENaCγ-VF mice were solubilized under non-denaturing conditions. γ-containing ENaC complexes were enriched by affinity capture via the C-terminal FLAG tag, eluted by 3C protease cleavage, and analyzed by mass spectrometry. **b.** Representative FSEC traces from lung, kidney, and colon showing mVenus fluorescence, confirming successful enrichment of γ-containing complexes. The gray shaded region indicates the fractions that were collected, pooled, and used for mass spectrometry analysis. A second prominent peak is observed and corresponds to background fluorescence detected in kidney tissue, independent of ENaC-containing complexes. **c.** Mass spectrometry results identifying proteins that co-purify with ENaCγ from each tissue. Data reveal both shared and tissue-enriched interactors, highlighting the power of this approach to define native ENaC-associated complexes in a tissue-specific context.

In addition to the core ENaC subunits, mass spectrometry revealed several regulatory proteins that associate with γ-containing assemblies. Notably, the E3 ubiquitin ligase Nedd4-2^43^ was detected across multiple tissues. Nedd4-2 is a well-established regulator of ENaC surface expression and turnover, mediating ubiquitin-dependent internalization and degradation. Its presence suggests that a subset of γ-containing complexes engages components of the ubiquitin regulatory pathway. We also detected the ER chaperone calnexin, consistent with its established role in the folding and quality control of multi-subunit membrane proteins^63–65^, including ENaC. Its presence suggests that another subset of γ-containing complexes engages components of the protein quality-control pathway. These associations support a model in which γ participates both in fully assembled ENaC channels and in regulatory-associated forms, providing a molecular basis for the broader elution profile observed by FSEC.

To evaluate the specificity of these interactions, we performed identical affinity purifications from wild-type tissues lacking the mVenus-3C-3xFLAG tag (Fig. S3). Neither Nedd4-2 nor calnexin was detected in these control samples, indicating that their presence in ENaCγ-VF purifications reflects specific co-purification with γ-containing complexes rather than nonspecific retention on the affinity resin. These findings broaden the scope of our analysis from subunit composition to the regulatory environment surrounding native ENaC. The ENaCγ-VF allele enables the isolation of both assembled αβγ channels, as well as γ-associated intermediary or regulatory complexes, linking biochemical heterogeneity observed by FSEC to defined molecular pathways that govern ENaC maturation, quality control, and turnover *in vivo*.

## Discussion

This study introduces the ENaCγ-VF knock-in mouse line as a physiologically faithful platform for dissecting the composition, assembly, and regulatory environment of native ENaC complexes across epithelial tissues. By combining endogenous fluorescent tagging with orthogonal biochemical and single-molecule approaches, this system overcomes longstanding challenges posed by ENaC’s low abundance, membrane localization, and complex subunit stoichiometry. Preservation of normal physiology in homozygous ENaCγ-VF mice demonstrates that the C-terminal mVenus-3C-3xFLAG fusion maintains native channel function, providing confidence that downstream biochemical and structural analyses reflect bona fide ENaC biology.

A notable strength of the ENaCγ-VF allele is its broad compatibility with both advanced and routine biochemical and imaging workflows. Here, we demonstrate that the mVenus tag enables sensitive detection by ensemble fluorescence, FSEC, confocal imaging, and TIRF microscopy. More generally, mVenus and FLAG tags also support robust detection by standard immunoblotting using well-validated commercial antibodies, making the mouse line readily compatible with widely-used biochemical analyses. In parallel, the C-terminal 3xFLAG epitope supports affinity purification followed by native-state elution using 3C protease. These features allow intact ENaC assemblies to be isolated without disrupting subunit composition or regulatory interactions. Thus, the ENaCγ-VF mouse provides a practical and adaptable foundation for examining ENaC at molecular, cellular, and tissue scales.

FSEC and SiMPull analyses reveal substantial heterogeneity among γ-containing ENaC complexes across lung, kidney, and colon. FSEC profiles identify both well-defined trimeric assemblies and broader, higher-molecular-weight populations, while SiMPull confirms that γ-containing complexes can be isolated and visualized at the single-molecule level under non-denaturing conditions. These findings indicate that ENaCγ participates in multiple molecular states *in vivo* and underscore the utility of the ENaCγ-VF system for resolving this complexity directly.

To distinguish fully assembled channels from intermediate or regulatory states, we developed 7B1-mScarlet, a fluorescent Fab that recognizes ENaCα under mild conditions. Dual-color FSEC showed that α-containing complexes elute within a narrower molecular range than γ-containing assemblies, suggesting preferential incorporation of α into a more restricted subset of αβγ channels. Validation in recombinant and native systems confirmed that 7B1-mScarlet selectively binds folded α within intact ENaC complexes.

Within this context, γ occupies a central position in the ENaC assembly landscape. Unlike α, which appears to be selectively incorporated into a narrower population of complexes, γ persists across multiple assembly and regulatory states, positioning it as a molecular hub for ENaC biogenesis and turnover. The broad elution range detected by FSEC, the presence of γ in both trimeric channels and regulatory-associated species, and its interaction with chaperones and ubiquitin ligases collectively suggest that γ participates in multiple checkpoints along the channel’s biogenesis, quality control, and turnover pathways. In contrast, α appears in a more restricted set of molecular contexts, consistent with its rate-limiting role in channel assembly and preferential incorporation into fully assembled αβγ complexes. The sharp, 7B1-positive peak therefore most likely reflects mature, fully assembled αβγ channels, whereas the broader γ profile encompasses assembly intermediates and regulator-engaged species. This interpretation is in line with quantitative biochemical studies showing an excess of β and γ over α subunits in native epithelia, implying that only a subset of γ participates in fully assembled channels at any given time^32^. These observations align with long-standing physiological models in which differences in subunit abundance govern ENaC assembly efficiency, but here are supported by direct biochemical analysis of intact native complexes. These findings indicate that native ENaC exists not as a single, static entity but as an ensemble of molecular species that differ in subunit composition, assembly state, and regulatory interactions. Decades of physiological and electrophysiological studies inferred such heterogeneity indirectly, but the lack of tools to visualize and isolate native complexes obscured the underlying molecular architecture^66^. The ENaCγ-VF model, particularly when combined with subunit-specific probes such as 7B1-mScarlet and native-state purification, begins to resolve this complexity with a level of precision that was previously inaccessible.

Mass spectrometry further supports this view by identifying regulatory factors that co-purify with γ-containing assemblies. In addition to α and β subunits, we detected Nedd4-2 and calnexin, key regulators of ENaC maturation, trafficking, and turnover. Their selective recovery in ENaCγ-VF purifications, demonstrates that affinity isolation via the γ subunit specifically enriches bona fide γ-associated ENaC complexes rather than nonspecific background proteins. The identification of Nedd4-2 and calnexin within these purifications further indicates that a subset of γ-containing assemblies engages quality-control machinery and regulatory pathways *in vivo*. This association provides a molecular explanation for the broader elution profile of γ-containing species and highlight the dynamic equilibrium between mature channels, assembly intermediates, and regulatory states *in vivo*. Taken together with the preserved physiology of ENaCγ-VF mice and the specific recovery of known ENaC regulators, these observations argue that the γ-containing species primarily reflect native assembly and quality-control states rather than artifacts of the γ-tag.

More broadly, the ENaCγ-VF platform enables a transition from indirect inference to direct molecular definition of native ENaC assemblies. By enabling isolation, visualization, and quantification of intact ENaC complexes under non-denaturing conditions, this system provides a foundation for linking subunit composition and regulatory state to tissue-specific ENaC function. As complementary tools for detecting additional ENaC subunits, particularly β, and for structural analysis of native complexes continue to emerge, the ENaCγ-VF model will serve as a foundational resource for defining the molecular principles that govern ENaC regulation across epithelial systems. In future work, this platform can be applied to examine how disease states remodel ENaC assembly and regulatory interactions, linking molecular architecture to disordered Na⁺ handling in conditions such as salt-sensitive hypertension, cystic fibrosis, and pseudohypoaldosteronism.

## Materials and Methods

### Generation of ENaCγ-VF mice

The ENaCγ-VF knock-in mouse line was generated by the Jackson Laboratory using CRISPR/Cas9-mediated genome editing to insert a sequence encoding mVenus, a 3C protease cleavage site, and a 3xFLAG tag at the C-terminus of the endogenous *Scnn1g* gene. Heterozygous animals (*ENaCγ-VF/+*) were initially shipped to our lab for validation. Following phenotypic and localization analysis (see Results), the Jackson Laboratory provided homozygous (*ENaCγ-VF/ENaCγ-VF*) mice, which were also viable and showed normal ENaCγ localization. Both heterozygous and homozygous mice are maintained under standard breeding protocols. To ensure continued integrity of the tagged allele, we routinely verify the line using genotyping services with primers designed to detect the presence of the mVenus-3xFLAG insertion. All animal procedures were conducted in accordance with protocols approved by the IACUC.

### Immunofluorescence Microscopy

Mice were anesthetized with a ketamine-xylazine-acepromazine cocktail (50:5:0.5 mg/kg). The kidneys were perfusion fixed by retrograde abdominal aortic perfusion of 3% paraformaldehyde in PBS (pH 7.4). After perfusion, the kidneys were removed, dissected, and cryopreserved in 800 mOsm/L sucrose in PBS overnight before being embedded in Tissue-Tek Optimal Cutting Temperature compound (Sakura Finetek, Torrance, CA). Slides were prepared by cutting 5 mm sections and stored at -80°C until use. For imaging, slides were incubated with 0.5% Triton X-100 in PBS for 30 minutes, blocked with 5% milk in PBS for 30 minutes, followed by incubation with primary antibody, diluted in blocking buffer, for 1 hour at room temperature or overnight at 4°C. Sections were then washed with PBS three times and incubated with fluorescent dye-conjugated secondary antibody, diluted in blocking buffer, for 1 hour at room temperature. Sections were washed with PBS three times and stained with 4’,6-diamidino-2-phenylindole before being mounted with ProLong Diamond Antifade Mountant (ThermoFisher Scientific, Carlsbad, CA). Images were captured using a KEYENCE BZ-X800 microscope (Itasca, IL).

### Amiloride treatment and urine analysis

Mice were injected intraperitoneally with vehicle (0.9% saline) and then placed in metabolic cages for a 6-hour urine collection. Five days later, the same animals received intraperitoneal injection of amiloride hydrochloride (40 μg per 25 g body weight), followed by another 6-hour urine collection. Sodium and potassium concentrations were measured by flame photometry, and the urinary Na⁺:K⁺ ratio was used to assess ENaC function.

### Filter binding assay

Kidney tissue was dissected from adult wild-type or ENaCγ-VF mice, rinsed in ice-cold tris buffered saline (TBS, 20mM Tris-HCl pH 7.6, 200mM NaCl), stored in TBS with protease inihibitors (Thermo Scientific™ Pierce protease inhibitor tablets, ThermoFisher Scientific) and flash-frozen in liquid nitrogen. For membrane preparation, frozen kidneys were pulverized using a liquid nitrogen-cooled mortar and pestle until a fine powder was obtained. Tissue was then homogenized in homogenization buffer (50mM phosphate buffer, pH 7.5) using a Dounce homogenizer. The homogenate was centrifuged at 1,800g for 10 min at 4 °C to remove nuclei and cellular debris. The resulting supernatant was further centrifuged at 100,000g for 60 min at 4 °C to pellet membrane fractions. Membrane pellets were resuspended in binding buffer (50mM phosphate buffer, pH 7.5). Equilibrium ligand binding was assessed using tritiated benzamil ([³H]-benzamil; Moravek). Membrane suspensions (typically 10 mg) were incubated with increasing concentrations of [³H]-benzamil (0.5-300 nM) in a final volume of 500 µL binding buffer. Incubations were carried out at 22 °C for 2 hours, a duration sufficient to reach equilibrium under these conditions. Nonspecific binding was determined in parallel samples containing 100 µM unlabeled phenamil (Phenamil mesylate, Tocris Bioscience). Binding reactions were terminated by rapid vacuum filtration onto presoaked glass microfiber filters (GF/B 25mm, Whatman) using a 12-well filtration manifold (Millipore). Filters were prewashed with 2ml ice-cold binding buffer and treated with 0.3% polyethyleneimine (PEI, MW 25K, Polysciences) to reduce nonspecific radioligand adsorption. Incubated samples were applied and each well was washed immediately with 2 x 5 mL ice-cold binding buffer. Filters were dried by vacuum, transferred to scintillation vials, extracted in 5 mL scintillation cocktail (Ultima Gold), and radioactivity was quantified by liquid scintillation counting (Beckman Coulter LS6500). Specific binding was calculated as total minus nonspecific counts per minute (CPM). Equilibrium binding curves were fitted to a one-site binding model using nonlinear regression in GraphPad Prism:

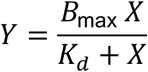

where *Y* is specific binding, *X* is the free ligand concentration, *B*ₘₐₓ is maximal binding, and *K_d_* is the apparent dissociation constant. Data from ENaCγ-VF and wild-type samples were fitted independently, and similar *K_d_* values were interpreted as evidence that the γ-tag does not perturb benzamil affinity.

### Tissue preparation and lysate generation for FSEC analysis

Tissues were harvested from ENaCγ-VF mice. For each FSEC experiment, one lung (all lobes), one kidney, and two colons were collected, rinsed in cold TBS, and weighed. Each tissue was homogenized in solubilization buffer containing 100 mM Tris-HCl (pH 7.6), 200 mM NaCl, 1% n-dodecyl-β-D-maltoside (DDM, Anagrade, Anatrace), protease inhibitors and Pefabloc® SC (Millipore Sigma). The homogenization volume was standardized to 750 µL per 100 mg of tissue. Samples were solubilized at 4 °C for 60 minutes with gentle rotation, then clarified by centrifugation at 100,000g twice for 40 minutes each. Supernatants were concentrated using a 100kDa centrifugal concentrator to a final volume of 110 - 480 µL. Concentrated samples were then centrifuged again at 100,000g for 10 minutes to remove aggregates. The final cleared supernatant was injected onto a Superose 6 Increase column (Cytiva) for FSEC analysis (Waters HPLC)^48^. Fractions were collected and analyzed based on mVenus fluorescence.

### SiMPull Assays

Glass coverslips were cleaned and PEGylated using standard silanization procedures^49^, then functionalized sequentially with streptavidin and a biotinylated anti-GFP/mVenus nanobody^67^. One experimental channel included the nanobody to capture fluorescently tagged ENaC subunits, while a parallel channel without nanobody was used to quantify background signal.

Streptavidin was applied at a concentration of 250 µg/mL and incubated for 5 minutes at room temperature, followed by a wash with the assay buffer (20 mM Tris-HCl (pH 7.6), 200 mM NaCl, 0.025% n-dodecyl-β-D-maltoside (DDM)) containing 200 µg/mL BSA. Biotinylated nanobody was then applied at 3 µg/mL and incubated for 10 minutes, followed again by a wash to remove unbound reagent. Clarified cell lysates (prepared as for FSEC analysis) containing fluorescently tagged ENaC subunits were then applied and incubated for 10 minutes at room temperature, followed by a final wash before imaging. Chambers were imaged on a Leica DMi8 TIRF microscope with a 100x oil-immersion objective. Images were captured using an Andor iXon Ultra 888 back-illuminated EMCCD camera with a 133 x 133 µm imaging area and 130 nm pixel size. For counting fluorophore spots, images were acquired using the excitation wavelength 488 nm. Spot detection and localization were performed using the ComDet plugin in ImageJ (FIJI). Molecule positions were analyzed within the center area of the field of view (512×512 px). The approximate particle size was set at 5 pixels. The spot intensity threshold (in SD) was set to 3. Data were statistically analyzed using GraphPad Prism.

### Affinity Isolation of Native ENaC Complexes for FSEC and Mass Spectrometry

Tissue preparation for this assay was performed using the same protocol as described for FSEC analysis, with the exception that digitonin was used as the detergent for solubilization instead of DDM. Briefly, lungs, kidneys, and colons were collected from 5 mice, and homogenized in solubilization buffer containing 100 mM Tris-HCl pH 7.6, 200 mM NaCl, 5mM EDTA, 1% digitonin (High purity, Millipore Sigma), protease inhibitors and Pefabloc. Clarified lysates were incubated with 50 µL of anti-FLAG M2 magnetic beads (Sigma-Aldrich) overnight at 4 °C with gentle rotation. Beads were washed 10 times with 15 ml wash buffer (20 mM Tris-HCl pH 7.6, 200 mM NaCl, 0.1% digitonin), and proteins were eluted by incubating with 25 µg of PreScission Proteases (GenScript) for 6 hours at 4 °C. The eluted material, in a total volume of 400 uL, was concentrated using centrifugal filters prior to injection onto a Superose 6 Increase 10/300 column (Cytiva) equilibrated in FSEC buffer (20 mM Tris-HCl pH 7.6, 200 mM NaCl, 0.1% digitonin). Fluorescence was monitored using an inline fluorimeter (Shimadzu RF-20A) to track mVenus-tagged ENaCγ. Fractions corresponding to the fluorescent signal were collected, pooled, and concentrated again for downstream mass spectrometry analysis.

### 7B1-mScarlet Fab Production

DNA sequences encoding the heavy and light chains of the Fab fragment of the anti-ENaCα monoclonal antibody 7B1 were cloned into separate pEG BacMam expression plasmids^68^. The heavy chain construct was engineered to include mScarlet-I at the C-terminus, followed by a Strep-tag II for affinity purification. Both constructs included an N-terminal leader sequence (MGWSCIILFLVATATGVHS) that allows for secretion of the Fab into the media. First passage of mBaculovirus was generated from each plasmid using Sf9 insect cells cultured at a density of 0.5×10⁶ cells/mL at 27 °C for 5 days. P1 virus was used to reinfect Sf9 insect cells cultured at a density of 1×10⁶ cells/mL at 27 °C for 4 days. After incubation, cells were pelleted by centrifugation at 1,800g for 20 minutes, and the supernatant containing the virus was harvested and filtered through a 0.22 µm filter. The resulting virus was used to infect suspension-adapted HEK293 cells at a density of 3×10⁶ cells/mL and incubated at 37 °C for 8 hours. After 8 hours, 10mM sodium butyrate was added and the cells were moved to a 30 °C incubator. Cells were cultured at 30 °C for 96 hours, after which they were spun down and the culture supernatant was collected and filtered again using a 0.22 µm filter. The clarified media with BioLock (1.6ml/L of FreeStyle™ 293, IBA Lifesciences) was loaded onto a 10 mL Strep-Tactin affinity resin (5ml/1L of media) (Cytiva) pre-equilibrated with wash buffer (20mM Tris pH 7.6, 200mM NaCl and 0.025% n-dodecyl-β-D-maltoside (DDM)). The column was washed with 20 column volumes of wash buffer, and bound 7B1-mScarlet Fab was eluted using desthiobiotin-containing wash buffer. Desthiobiotin (IBA Lifesciences) was subsequently removed using a PD-10 desalting column (Cytiva), and the Fab was eluted in wash buffer. Purity and labeling of the Fab were confirmed by SDS-PAGE and FSEC.

### Cell-surface biotinylation and validation of 7B1-mScarlet binding to recombinant ENaC

To evaluate binding of the 7B1-mScarlet Fab to recombinant mouse ENaC, we expressed a mouse ENaC construct composed of the α (Uniprot ID Q61180), β (Uniprot ID Q9WU38), and γ (Uniprot ID Q9WU39) subunits, with the γ subunit bearing the C-terminal mVenus tag. Cells were analyzed under both total and plasma membrane-enriched conditions. For total protein analysis, cells were solubilized directly^36, 37^, clarified by centrifugation, and the resulting lysates were incubated with 7B1-mScarlet prior to downstream analysis. To specifically assess plasma membrane-expressed ENaC, cell-surface proteins were labeled using a membrane-impermeant biotinylation reagent according to the manufacturer’s instructions (Pierce™ Cell surface biotinylation & Isolation Kit, ThermoFisher Scientific). Briefly, cells expressing recombinant mouse ENaC were harvested and incubated with biotin (EZ-Link Sulfo-NHS-SS-Biotin) to selectively label surface-exposed proteins. Excess biotin was quenched and removed by washing, after which cells were either flash-frozen for storage at -80 °C or processed immediately for solubilization.

Cells were lysed in buffer containing 20 mM Tris-HCl (pH 7.6), 150 mM NaCl, 20mM DDM, 3mM CHS, and protease inhibitors. Lysates were clarified by centrifugation, and the supernatant was incubated with GFP nanobody resin (GNB). Following binding, the resin was washed with 20mM Tris (pH7.6), 150mM NaCl, 0.5mM DDM, 75uM CHS without and with 5mM CaCl2. Both biotinylated and non-biotinylated fractions of the protein were eluted by thrombin (30ug/ml of resin, Human alpha-thrombin, Prolytix) cleavage. The thrombin eluted sample was passed through the Strep-Tactin resin to capture biotinylated, plasma membrane-expressed proteins. The unbound flow-through, representing the intracellular protein fraction, was collected. Following fractionation, 7B1-mScarlet was added to the flow-through fraction. Biotinylated proteins bound to the Strep-Tactin resin were subsequently eluted using excess biotin (10x Buffer BXT, IBA Lifesciences), and 7B1-mScarlet was added to the eluted plasma membrane fraction. Both intracellular (flow-through) and plasma membrane (biotin-eluted) fractions were analyzed by HPLC, with fluorescence monitored in the mScarlet channel.

## Data Availability Statement

All data in this manuscript are available from the corresponding author upon request.

## Acknowledgements

We thank members of the Gouaux laboratory, especially Eric Gouaux, April Goehring, and Natalie Sheldon, for their generous guidance and support as we established animal-based experiments and native-state biochemical workflows. We are grateful to Ashok Reddy and the staff of the OHSU Proteomics Shared Resource for mass spectrometry analyses and technical expertise. We thank the OHSU Department of Comparative Medicine for assistance with animal care and colony management. The ENaCγ-VF mouse line was generated by The Jackson Laboratory. Funding for this project was provided by institutional support from the Vollum Institute and by NIH grant R01GM138862 to I.B, R01DK132066 to J.A.M, and R01s DK133220 and DK51496 to D.E.

## Author Contributions

A.B. maintained the mouse colony, harvested mouse tissues, and performed FSEC, SiMPull, and purification of native ENaC complexes. J.C. performed the amiloride challenge experiments. X.-T.S. conducted confocal imaging of heterozygous and homozygous mice. R.B.M. maintained the mouse colony and harvested mouse tissues. J.A.M. and D.H.E. supervised the physiology experiments. I.B. conceived and supervised the study. All authors contributed to manuscript preparation.

---

**Figure S1. Immunofluorescence staining of kidney tissue from heterozygous ENaCγ-VF mice.**

**a**. Representative immunofluorescence image showing co-localization of mVenus, ENaCγ, and aquaporin-2 in kidney tissue. mVenus signal was detected using a FITC-conjugated anti-GFP primary antibody (goat, Rockland, 1:100) and Alexa Fluor 488 donkey anti-goat secondary antibody. Endogenous ENaCγ was detected using a rabbit anti-ENaCγ primary antibody (StressMarq, 1:500) targeting the C-terminus, and Alexa Fluor 647 donkey anti-rabbit secondary antibody. Aquaporin-2 (AQP2) was stained using a mouse anti-AQP2 primary antibody (Santa Cruz, 1:50) and Alexa Fluor 555 donkey anti-mouse secondary antibody. **b**. Representative immunofluorescence image showing co-localization of mVenus, ENaCα, and pNCC in kidney tissue. mVenus signal was detected using a FITC-conjugated anti-GFP primary antibody (goat, Rockland, 1:100) and Alexa Fluor 488 donkey anti-goat secondary antibody. Endogenous ENaCα was detected using a rabbit anti-ENaCα primary antibody (generous gift of J. Loffing^69^) targeting the C-terminus, and Alexa Fluor 647 donkey anti-rabbit secondary antibody. The phosphorylated form of the Na⁺-Cl⁻ cotransporter (NCC) was stained using a chicken anti-NCC (generously provided by J. Wade^70^) and Cy3 donkey anti-chicken secondary antibody. **c**. Close-up view highlighting localization of tagged mVenus, ENaCα, and pNCC. * indicate DCT1 (pNCC+/ENaCα-); + indicate DCT2 (pNCC+/ENaCα+).

**Figure S2. Validation of 7B1-mScarlet as a tool for detecting mouse ENaCα.**

**a.** Schematic of the recombinant mouse ENaC subunit constructs used for validation. The α and β subunits were expressed with wild-type sequences, while the γ subunit was C-terminally fused to mVenus, mirroring the configuration in ENaCγ-VF mice. **b, c.** FSEC traces showing mVenus (b) and mScarlet fluorescence (c) from lysates of cells expressing recombinant mouse ENaC. Co-elution of the two signals confirms that 7B1-mScarlet binds specifically to complexes containing ENaCα, validating its use in detecting α subunits within γ-containing assemblies. **d.** Schematic illustration of ENaC complexes located inside the cell and in the plasma membrane. Only surface ENaC is exposed to membrane-impermeable biotin. **e.** FSEC traces of purified surface-expressed ENaC protein (black trace) and intracellular ENaC pool (blue trace) monitored on the mScarlet channel fused to the 7B1 Fab. The difference in peak height is due to preexisting differences in relative abundance and not a difference in 7B1 recognition.

**Figure S3. Control purification workflow using wild-type mice to assess non-specific binding.**

Schematic of the purification workflow applied to wild-type mouse tissues, performed in parallel with the ENaCγ-VF workflow. Although the protocol is identical, including tissue solubilization, anti-FLAG affinity capture, and protease elution, wild-type mice lack the C-terminal 3xFLAG tag on ENaCγ and therefore cannot be captured by the anti-FLAG resin. This control serves to identify proteins that may non-specifically bind to the affinity resin and distinguishes true ENaC-associated interactors from background contaminants in the ENaCγ-VF purification experiments.

