## Supplementary figures and images for "Differential Assembly of Native ENaC Complexes Across Mouse Epithelial Tissues"

### Supplemental Figure

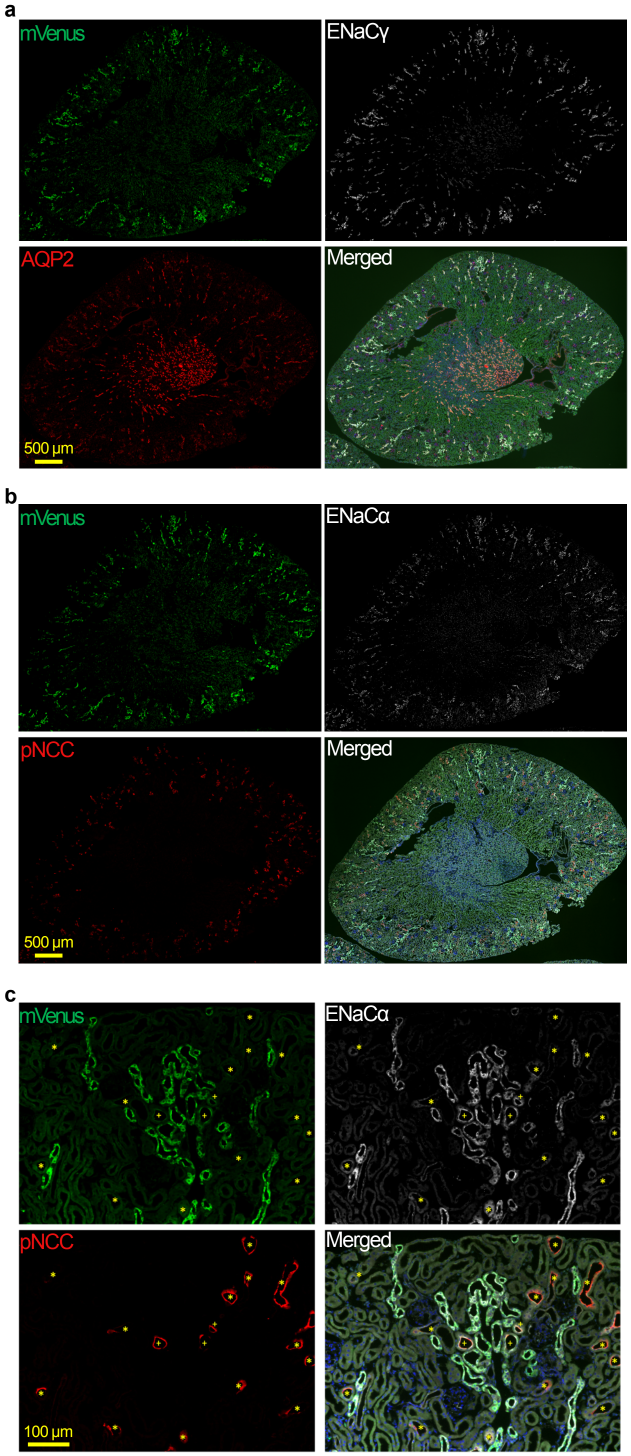

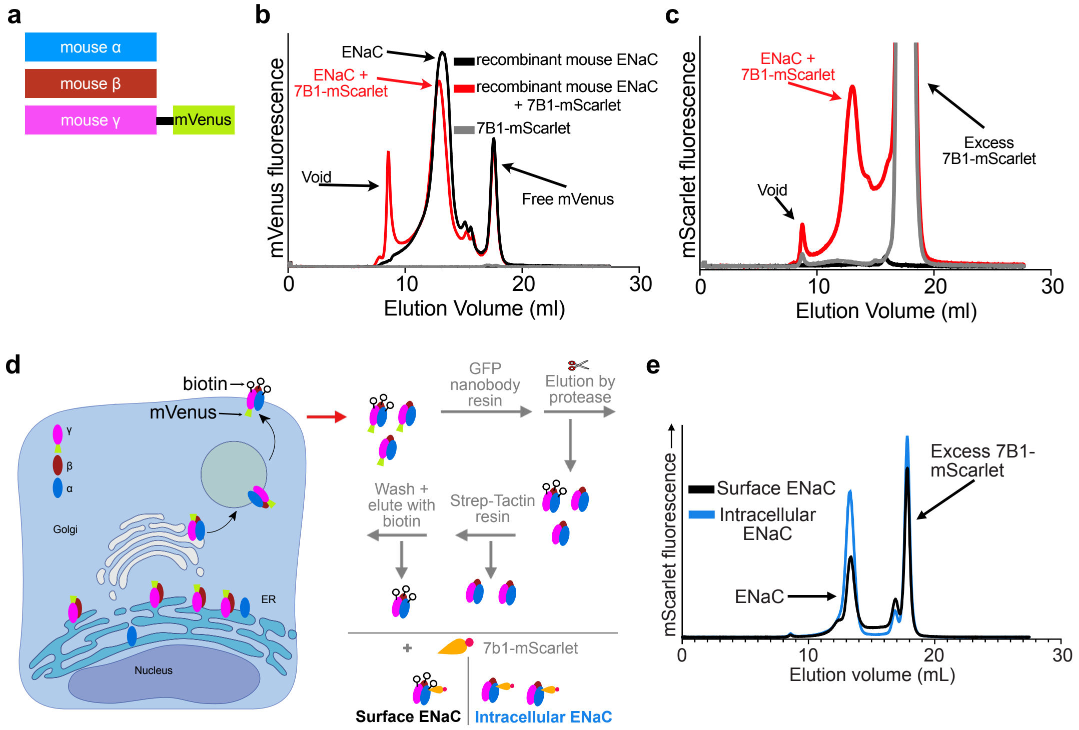

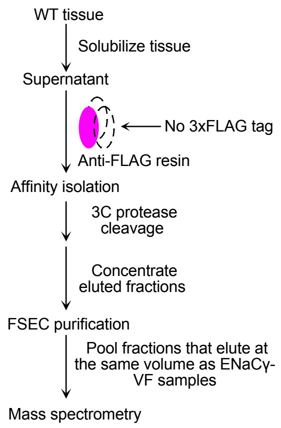
